# Off-the-Shelf Multilayer Vascular Grafts with Damage-Resistant Hydrogel Coatings Incorporating Integrin Targeting

**DOI:** 10.64898/2026.07.28.741222

**Authors:** Abbey Nkansah, Ashauntee Fairley, Nicolai Ang, Maddie Laude, Andrew Robinson, Nicholas Grammer, Xiaoxiao Zhang, Lee-Jae Jack Guo, Mir Timo Zadegh Nazari Shafti, Abdelmotagaly Elgalad, Elizabeth Cosgriff-Hernandez

## Abstract

Synthetic grafts remain ineffective for small-caliber vascular applications due to thrombosis and intimal hyperplasia. To address these limitations, our lab designed a multilayer graft consisting of a hydrogel coating that promotes post-implantation endothelialization and an electrospun mesh that matches arterial mechanical properties. Damage-resistant hydrogels were engineered using a double-network system composed of polyether urethane diacrylamide and N-acryloyl glycinamide to enhance fracture resistance through hydrogen bonding. In this study, we utilized redox initiation to apply conformal, durable hydrogels to electrospun grafts. Bioactivity was introduced using streptococcal collagen-like proteins containing α1β1 and α2β1 integrin-binding motifs, enabling selective cell–material interactions that support endothelialization while preserving acute thromboresistance. To establish the feasibility of these grafts as off-the-shelf devices, we evaluated coating integrity and bioactivity retention following sterilization and dynamic physiological loading. Sterilized composites exhibited surgically-associated damage resistance, indicating that sterilization did not compromise hydrogel durability. Coating integrity and bioactivity were also preserved after six weeks of physiological loading. Acut thromboresistance was supported by both static platelet adhesion assays and dynamic whole-blood bioreactor studies using heparinized blood, with low platelet adhesion observed relative to ePTFE. Finally, a pilot ovine carotid model demonstrated successful surgical handling and sustained graft patency. Collectively, these results highlight the promise of multilayer vascular grafts as durable, thromboresistant conduits for small-diameter vascular applications.

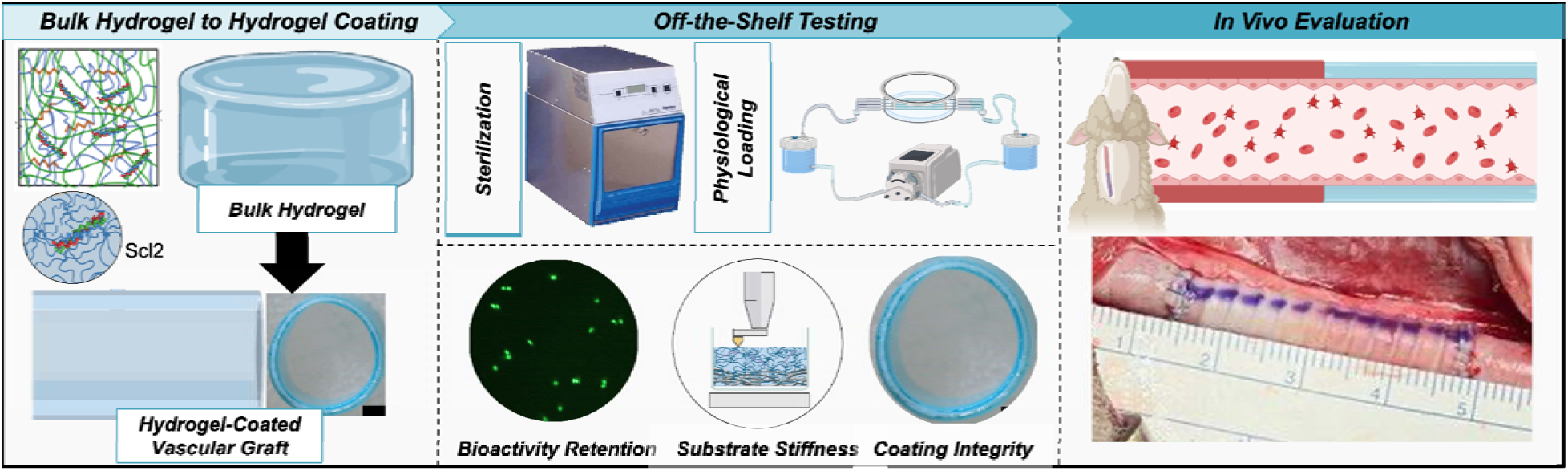

## Introduction

Cardiovascular disease is the leading cause of death in the United States with coronary artery and peripheral artery disease contributing to more than 25 million cases annually.^1^ Of these cases, nearly 400,000 patients in the United States require bypass graft surgeries each year to redirect blood flow around vessels occluded from extensive thrombus formation and smooth muscle cell hyperproliferation.^1–3^ Autologous vessels are the current gold standard as replacements for arterial prostheses but are often unavailable due to size mismatch and multiple cases of vascular trauma.^2, 4^ Synthetic vessel replacements as supplements to autologous vessels during multi-grafting procedures are highly desirable to increase treatment options for patients with severe cases of coronary artery and peripheral vascular disease.^5, 6^ However, thrombogenicity and low compliance of current synthetic graft materials (e.g., Dacron®, Gore-Tex®) limits their efficacy in small diameter applications.^7–9^ These failure modes arise from the absence of a quiescent endothelial cell monolayer and poor matching of arterial compliance.^10–12^ Therefore, there is a critical need to develop alternative off-the-shelf, synthetic vascular graft that can function in small diameter applications.

To address these disparate biological and mechanical failure mechanisms, our lab developed a multilayer vascular graft comprised of an electrospun mesh that matches the arterial mechanical properties coated with a thromboresistant, bioactive hydrogel to provide luminal endothelialization.^13, 14^ The hydrogel coating is based on a polyethylene glycol (PEG) macromer with a collagen-mimetic protein derived from group A Streptococcus, Scl2.28 (Scl2).^15, 16^ The streptococcal collagen-like proteins contain α1β1 and α2β1 integrin-binding motifs, enabling selective cell–material interactions that support endothelialization while preserving acute thromboresistance.^15, 16^ This graft provides an alternative strategy to current endothelialization approaches that require pre-seeding and extended culture periods to achieve endothelialization, precluding their use in emergency procedures and incurring higher costs.^17–20^ However, the brittle nature of the PEG-based hydrogel prevented its surgical use as particulate generation from suturing and hydrogel layer damage was observed when deployed as interposition artery graft in a pig model.^21^ To address this issue, a damage-resistant interpenetrating network (IPN) hydrogel chemistry was developed with high extensibility and defect tolerance required for durability while maintaining facile protein incorporation and substrate stiffness required for cell attachment.^13, 22^ This new hydrogel formulation has shown great potential in mitigating damage during suturing, torquing, and stretching during implantation.^13^ Extending these formulations from simplified two-dimensional substrates to fully three-dimensional grafts introduce additional design considerations to ensure retained conformability and bioactivity retention. Specifically, additional testing is needed to confirm that findings from previous two-dimensional substrate studies are maintained when the hydrogel is applied as a coating in the multilayer graft.^13, 23^ For example, we have previously reported the effect of hydrogel modulus on endothelial cell attachment.^23^ Therefore, it is important to evaluate whether the cells interacting with the multilayer grafts sense the underlying electrospun mesh as these differences may introduce heterogeneous cell mechanosensing that can negatively modulate integrin-mediated endothelial responses.^24^ In addition, utility as an off-the-shelf vascular graft requires that the hydrogel coating display uniformity and stability after drying, sterilization, and prolonged physiological loading. Bioactivity retention of the integrin-targeting hydrogel for long periods under physiological flow is also necessary to ensure sustained ligand presentation throughout the endothelialization process.

Herein, we report on the fabrication of a multilayer vascular graft using this newly developed durable IPN hydrogel and assessed its suitability as an off-the-shelf synthetic conduit for bypass and other vascular interventions. First, bulk IPN hydrogels were compared to multilayer vascular grafts to ensure that damage resistance and bioactivity were retained when the bioactive hydrogel was applied as a conformal graft coating. Multilayer vascular grafts were then dried and sterilized with ethylene oxide to confirm retention of bioactivity and damage resistance when processed as an off-the-shelf graft. Assessments of thromboresistance using benchtop static platelet attachment and a dynamic whole blood bioreactor was utilized to establish resistance to platelet attachment of the multilayer graft. Benchtop testing of multilayer grafts under physiologically relevant loading conditions was performed for up to 6 weeks to confirm hydrogel coating integrity and bioactivity retention prior to pilot in vivo evaluation in an ovine carotid interposition graft model. Collectively, these studies were used to assess the potential of multilayer vascular grafts with durable, bioactive coatings to function as off-the-shelf small-diameter conduits.

## Results

Towards translation of bulk hydrogel slabs to coatings for vascular graft applications, it is critical to establish comparable mechanical and biological properties to ensure accurate translation of these simple hydrogel slab studies to multilayer graft performance (**Figure 1A)**. Hydrogel slab stiffness was evaluated in comparison to the stiffness of the hydrogel coating to determine difference between multilayer grafts and prior cell studies on hydrogel specimens (**Figure 1B)**.^13^ Similarly, bioactivity retention was assessed by quantifying protein incorporation, cell adhesion, and cell spreading after multilayer vascular graft fabrication in comparison to bulk hydrogel slabs. Similar levels of Scl2_GFPGER_ incorporation were observed between bulk gels and the hydrogel coating (**Figure 1C)**. As expected given the similar protein incorporation, there were no significant differences in endothelial cell adhesion and spreading between bulk hydrogel slabs and hydrogel coatings (**Figure 1D, E)**. Collectively, this indicates that the cells respond solely to the hydrogel coating properties with no observable effect of the underlying electrospun mesh.

**Figure 1:**
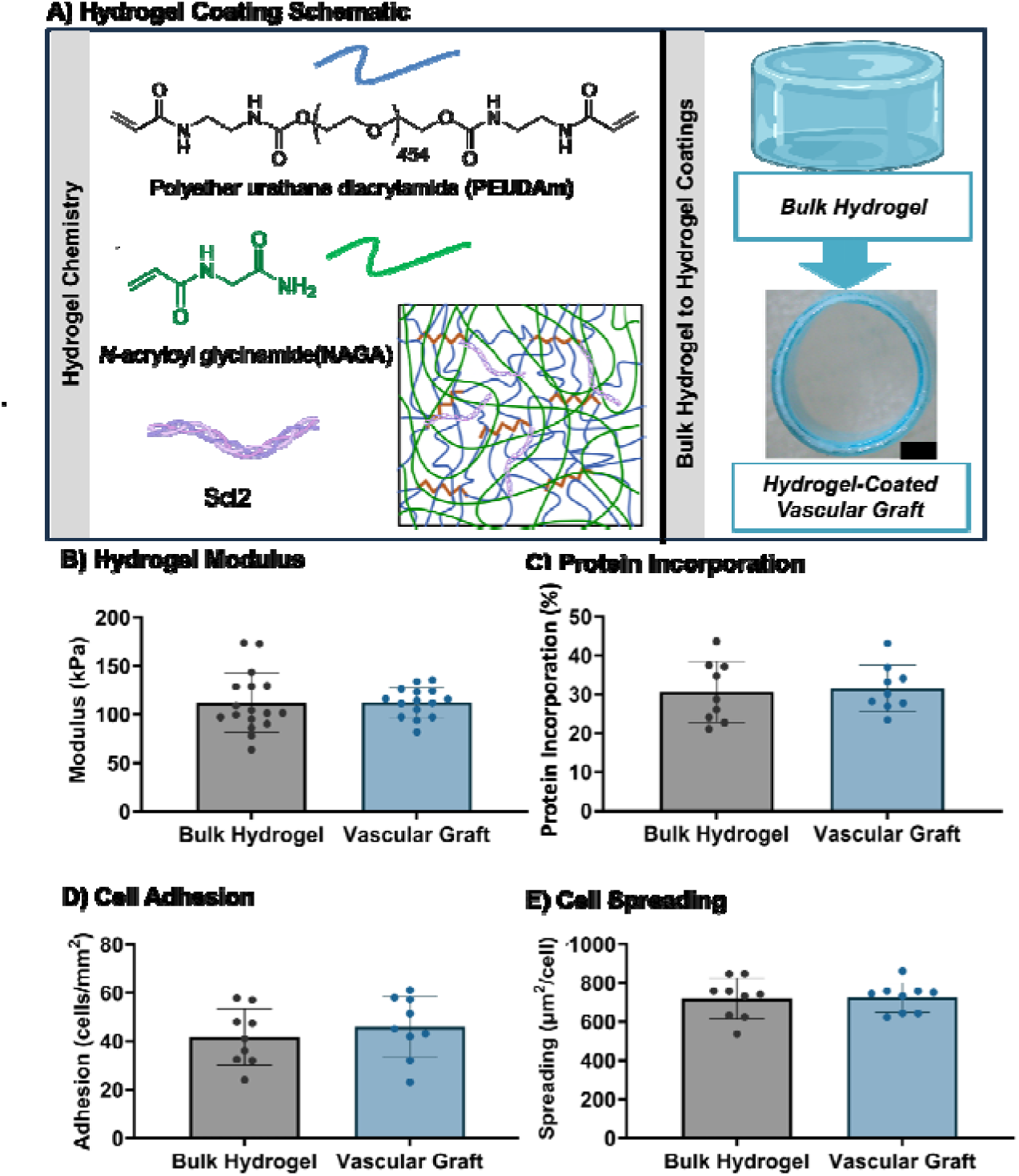
Effect of hydrogel form factor (hydrogel slab vs hydrogel coating) on modulus, protein incorporation, and bioactivity. A) Schematic representation of hydrogel chemistry and translation of hydrogel slabs to hydrogel coatings (scale bar = 1 mm) B) Hydrogel modulus (n = 3 grafts, 4 indentations per graft). C) Scl2 protein content (n = 3 specimens per graft, 3 grafts). Endothelial cell adhesion (n = 3 specimens per graft, 3 grafts). E) Cell spreading (n = 3 specimens per graft, 3 grafts). Data for all comparisons represent the average and standard deviation of each specimen. Comparisons reported showed no statistically significant differences.

### Effect of drying and sterilization on bioactivity retention

To demonstrate utility as an off-the-shelf medical device, multilayer vascular grafts must support endothelialization following drying and sterilization. The multilayer vascular graft demonstrated no statistically significant differences in modulus following drying and sterilization in comparison to control specimens (**Figure 2B)**. As expected, protein retention was unaffected following drying and sterilization (**Figure 2C)**. Cell adhesion and spreading studies revealed that sterilization had minimal impact on bioactivity retention on multilayer vascular graft sections, as noted by no statistically significant differences in cell adhesion and spreading (**Figure 2D, E)**.

**Figure 2:**
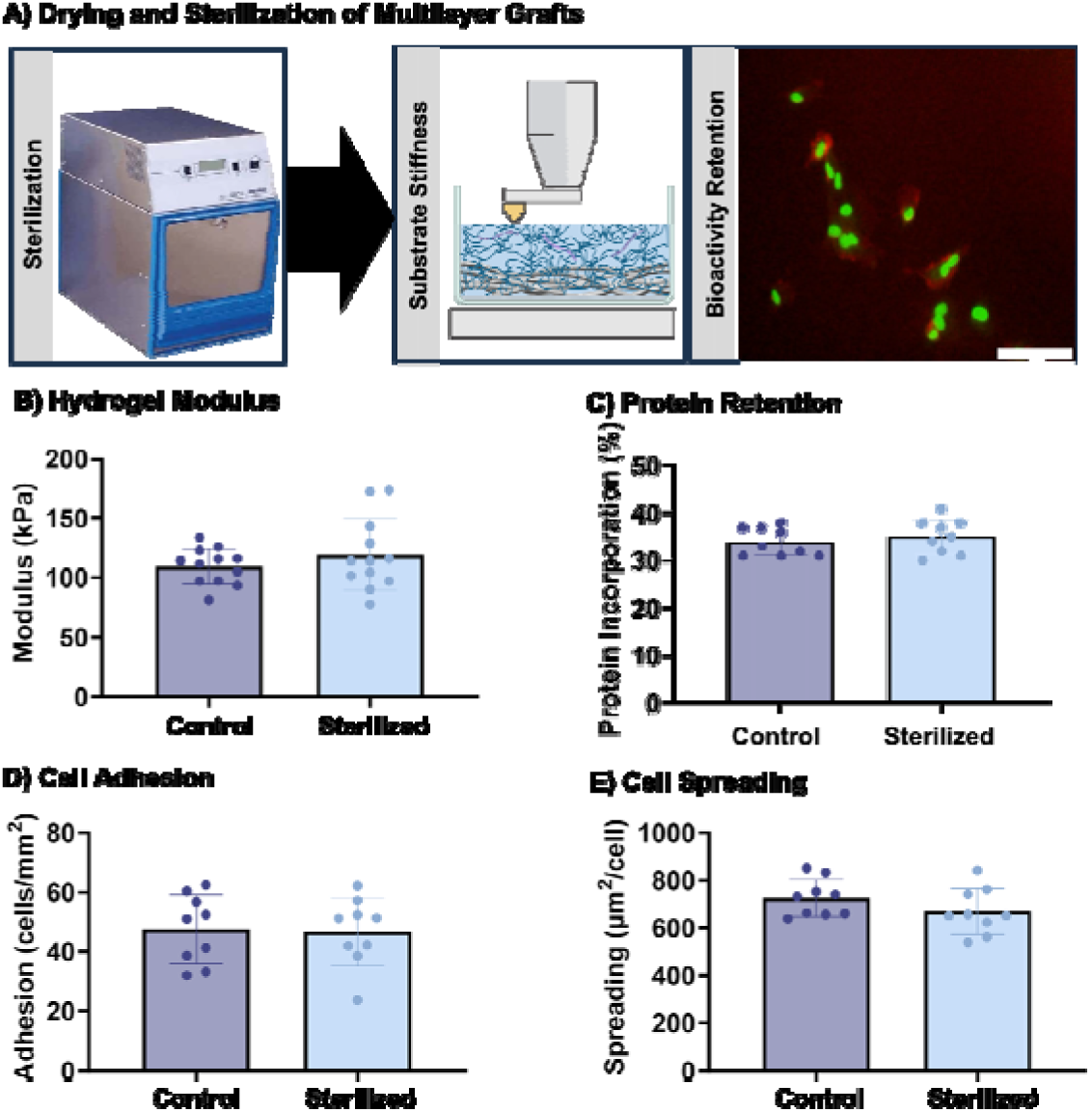
Characterization of multilayer vascular graft bioactivity retention following drying and sterilization. A) Schematic of multilayer graft drying and ethylene oxide sterilization process B) Hydrogel modulus (n = 3 grafts, 4 indentations per graft). C) Scl2 protein content (n = 3 specimens per graft, 3 grafts). D) Endothelial cell adhesion (n = 3 specimens per graft, 3 grafts). Cell spreading (n = 3 specimens per graft, 3 grafts). Data for all comparisons represent the average and standard deviation of each specimen. Comparisons reported showed no statistically significant differences.

These studies indicate that drying and sterilization had minimal impact on the bioactivity retention of the multilayer graft.

### Effect of drying and sterilization on hydrogel damage resistance

Mitigation of damage to the hydrogel coating is crucial for surgical implantation, as damage to the hydrogel layer can pose risks of thrombus formation and embolism.^21^ Therefore, we characterized multilayer graft damage resistance after drying and sterilization procedures in comparison to a positive control, a brittle PEGDA 3.4 kDa hydrogel coating.^13, 21^ Following sterilization, there was no statistical difference in particulate generation with suturing following drying and sterilization of IPN composites (**Figure 3A)** and significantly fewer dislodged particles as compared to the PEGDA hydrogel control. Resistance to torque-induced damage was also evaluated on sections of multilayer grafts following sterilization. Compared to the PEGDA hydrogel graft controls which showed visible signs of damage following torquing the grafts 180°, IPN coated vascular grafts exhibited no visible signs of torque-induced damage following torquing before or after drying and sterilization (**Figure 3B)**. Stretching damage on sectioned vascular graft samples was used as a qualitative assessment of cohesive strength retention following sterilization. The control PEG hydrogel coating displayed hydrogel coating damage prior to reaching 100% strain for all specimens; whereas, none of the IPN multilayer grafts displayed visible signs of damage to the hydrogel layer following sterilization (**Figure 3C)**. Damage resistance of the IPN multilayer grafts was attributed to high extensibility, tensile strength, and fracture energy of the IPN hydrogel formulation.^25^ These results indicate that damage resistance of the IPN hydrogel coating was retained after drying and sterilization.

**Figure 3:**
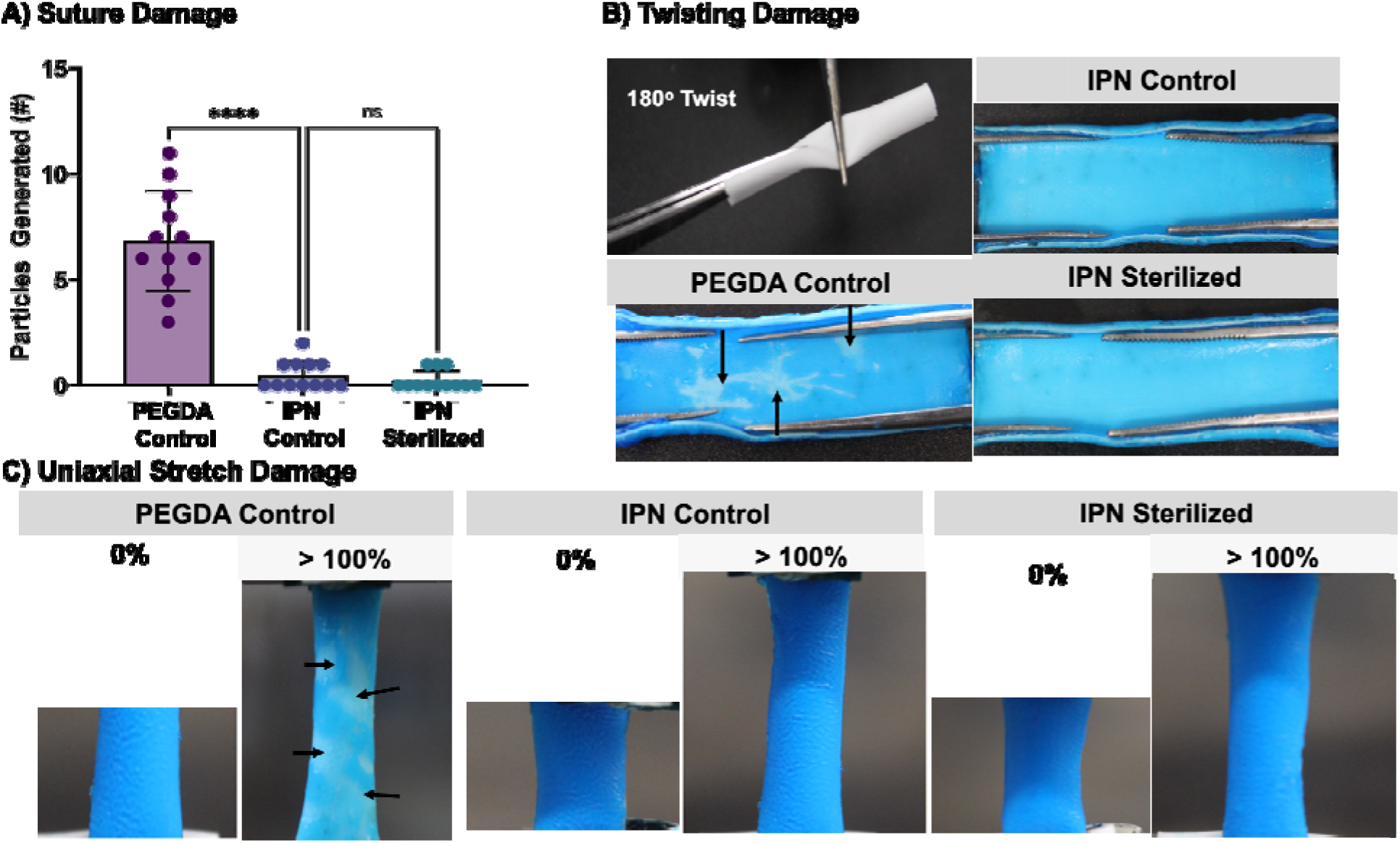
Effect of drying and sterilization on damage resistance of IPN hydrogel multilayer grafts in comparison to PEGDA hydrogel multilayer graft control. A) Suture damage to hydrogel coating and particle generation during suturing (n = 4 per graft, 3 grafts). B) Damage of hydrogel coating on tubular constructs after gripping and twisting with forceps (n = 3 per grafts per condition) (arrows signify damaged regions). C) Damage of hydrogel coatings after uniaxial stretch (n = 3 grafts per condition) (arrows signify damaged regions). Mean and standard deviation represented for suture damage characterization (ns = not significant; **** = p < 0.0001). Full image data sets are provided in the data repository.

### Bioactivity retention following physiological loading

A custom flow loop was used to determine the effect of physiological conditioning on the multilayer grafts after 24 h, 1 week, or 6 weeks of loading at physiological temperature (**Figure 4A)**. Multilayer grafts maintained hydrogel coating integrity and patency after extended physiological loading with no visible signs of damage following 24 h, 1 week, and 6 weeks (**Figure 4B)**. Hydrogel stiffness, protein concentration, and bioactivity after this extended conditioning was compared to static incubation and the untreated control. Nanoindentation of multilayer grafts indicated that there were no significant differences in moduli between physiologically loaded hydrogel coatings and the static and untreated controls at any time point (**Figure 4C).** Protein retention was similarly maintained following physiological loading as both unloaded and loaded grafts showed comparable levels of Scl2 protein retention, with no statistically significant reductions after 6 weeks of physiological conditioning (**Figure 4D)**. The constructs exposed to static and dynamic flow conditions for 6 weeks also displayed similar levels of cell attachment for 6 weeks in comparison to earlier time points, indicating that the protein maintained targeted integrin interactions over this period (**Figure 4E)**. These studies confirm that there was minimal loss of protein and bioactivity from the multilayer grafts for extended periods under physiological loading.

**Figure 4:**
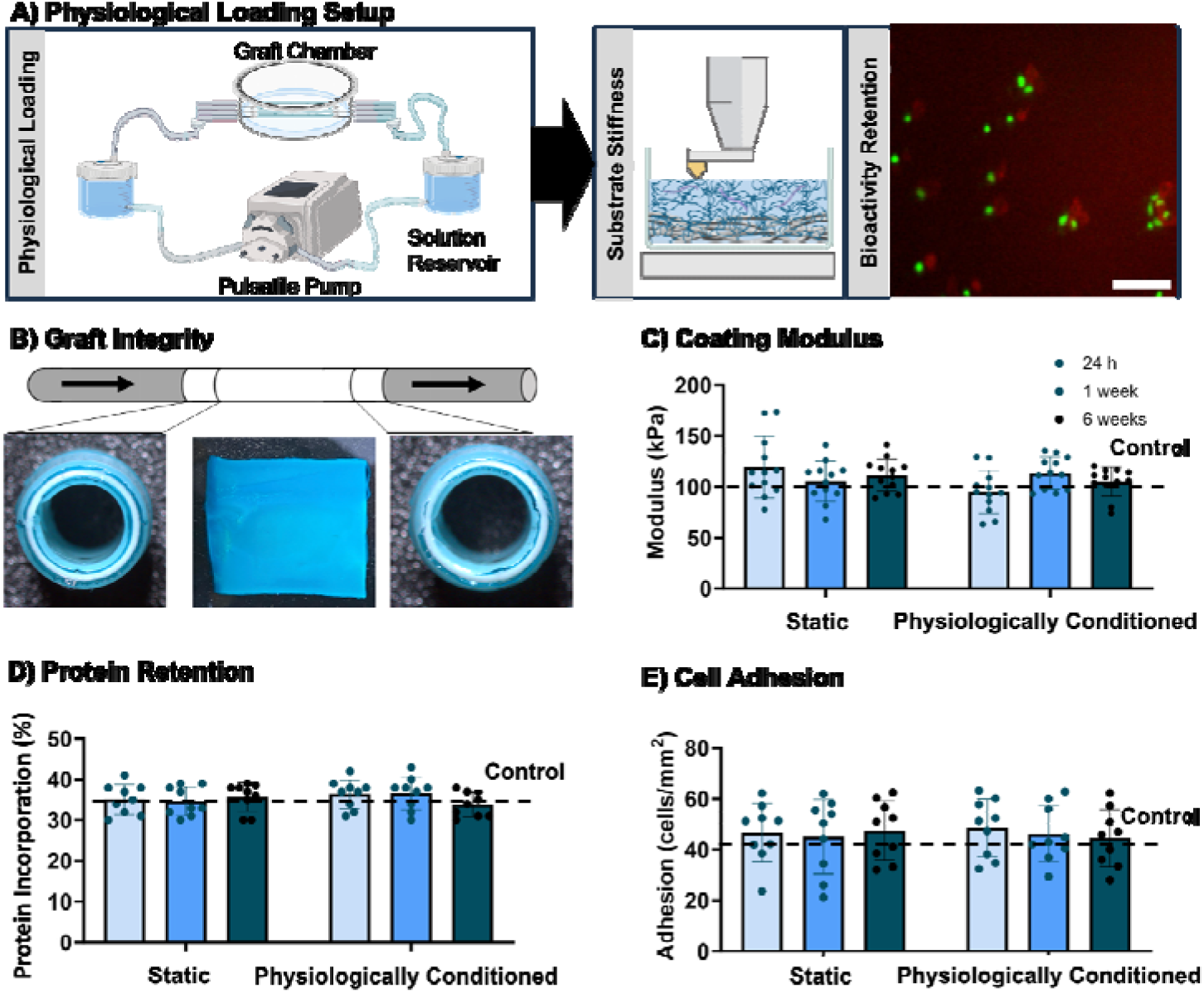
The effect of physiological loading on hydrogel coating integrity and bioactivity retention over time. A) Schematic of physiological loading setup with assessments performed at 24 h, 1 week, and 6 weeks B) Resistance of sterilized IPN hydrogel coated tubular grafts to delamination after 6 weeks of pulsatile flow. C) Hydrogel modulus (n = 3 grafts, 4 indentations per graft). D) Protein quantitation of multilayer grafts (n = 3 specimens per graft, 3 grafts per condition). E) Endothelial cell adhesion (n = 3 specimens per graft, 3 grafts per condition). Data for all comparisons represents the average and standard deviation for each condition.

### Acute thromboresistance assessment of multilayer graft

As a preliminary assessment of acute thromboresistance, platelet attachment to the multilayer grafts was assessed under static exposure and dynamic testing using a whole blood bioreactor under arterial-like flow in comparison to an ePTFE clinical control graft. Platelet attachment was significantly reduced on multilayer grafts as compared to ePTFE grafts under static conditions (**Figure 5A)**. Multilayer grafts also withstood occlusion due to clotting for the entirety of the 6-hr flow study **(Figure 5B)**. In contrast, the ePTFE graft occluded within 1-2 hours due to substantial clot formation. Decreased levels of platelet attachment were also evident on multilayer grafts as compared to the ePTFE graft. These results support the development of a thromboresistant multilayer graft with reduced platelet attachment with direct comparison to current clinically available graft alternative.

**Figure 5:**
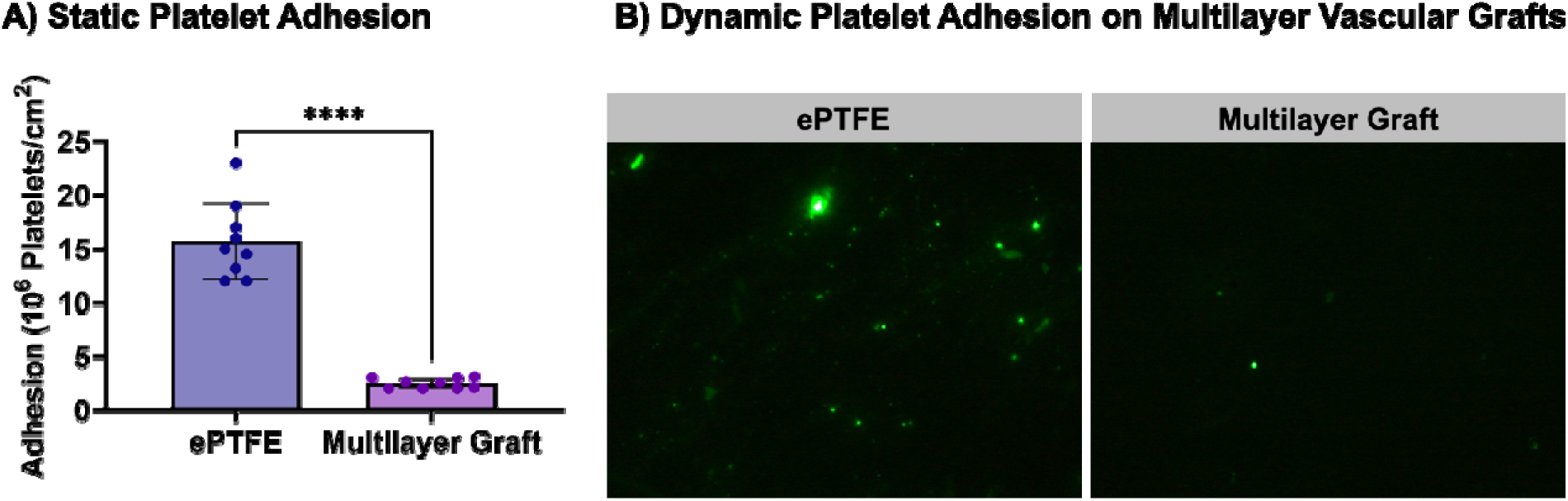
In vitro characterization of acute thromboresistance. A) Static platelet attachment multilayer grafts and ePTFE (GORE-TEX®, Gore Medical) vascular grafts. B) Images of meparcine-labeled platelets after 6h of whole blood flow in a pulsatile flow bioreactor (n = 3 grafts, 3 specimens per graft). Mean and standard deviation represented for platelet adhesion characterization (**** = p < 0.00001).

### Pilot assessment of multilayer graft in an ovine model

To investigate surgical and *in vivo* performance of this multilayer graft, coil-reinforced multilayer grafts were implanted in an ovine carotid artery interposition graft model with a target duration of 4 weeks *in vivo* (**Figure 6A**). Coil-reinforced vascular grafts were chosen for *in vivo* studies to provide the requisite kink resistance to prevent graft occlusion.^14^ To ensure arterial size matching, sheep underwent preoperative imaging to estimate carotid artery diameter (5.3 – 5.7 mm ID) and multilayer grafts were fabricated in comparable size ranges (5.0, 5.5, 6.0 mm ID) and selected based on each surgical sheep’s carotid artery sizing. Across all cases, successful anastomoses were performed with hemostasis established and grafts maintaining pressure with no signs of fluid leakage or kinking. Acute thromboresistance was established as indicated by doppler ultrasonography displaying no occlusion; however, only one of the cases was monitored for the planned 30 days. The first two animal studies provided critical information regarding optimization of postoperative management and longitudinal monitoring protocols. In the first case, the postoperative anticoagulation regimen consisted of aspirin, clopidogrel, heparin, and warfarin, with Doppler ultrasound surveillance performed weekly. The graft remained patent on day 7; however, by day 14 intraluminal thrombosis was detected, resulting in graft occlusion. Based on serial postoperative imaging demonstrating thrombus progression, the postoperative protocol was revised to include more intensive anticoagulation management with more frequent coagulation monitoring and Doppler ultrasound surveillance to enable earlier detection of thrombus formation and timely adjustment of anticoagulation therapy. In the second case, the revised postoperative anticoagulation and monitoring protocol was implemented, and the graft demonstrated sustained thromboresistance during the observation period. However, the animal was terminated prematurely on postoperative day 3 due to reduced food intake unrelated to the graft or surgical procedure, with no evidence of stroke, pulmonary embolism, cardiac events, or peripheral thrombus formation. The optimized postoperative management strategy was continued in the third case, which maintained graft patency through the full 30-day study period. No significant intraluminal thrombosis was observed before the postoperative day 25 and patency was maintained for the full study (**Figure 6B**). Intermittent intraluminal faint thrombi were noted during the final five days, which was resolved by increasing the anticoagulation regimen. All multilayer grafts displayed intact hydrogel coating integrity with no signs of degradation or delamination from the underlying substrate following explantation, which was the primary outcome assessment of this pilot study. Future *in vivo* studies will incorporate additional replicates with this revised protocol to further validate the long-term results. These findings highlight the importance of iterative refinement of animal protocols when evaluating novel blood-contacting biomaterials and provide important considerations for future validation studies. Additional replicates using the optimized protocol will be required to determine the long-term thromboresistance and patency of the multilayer graft.

**Figure 6:**
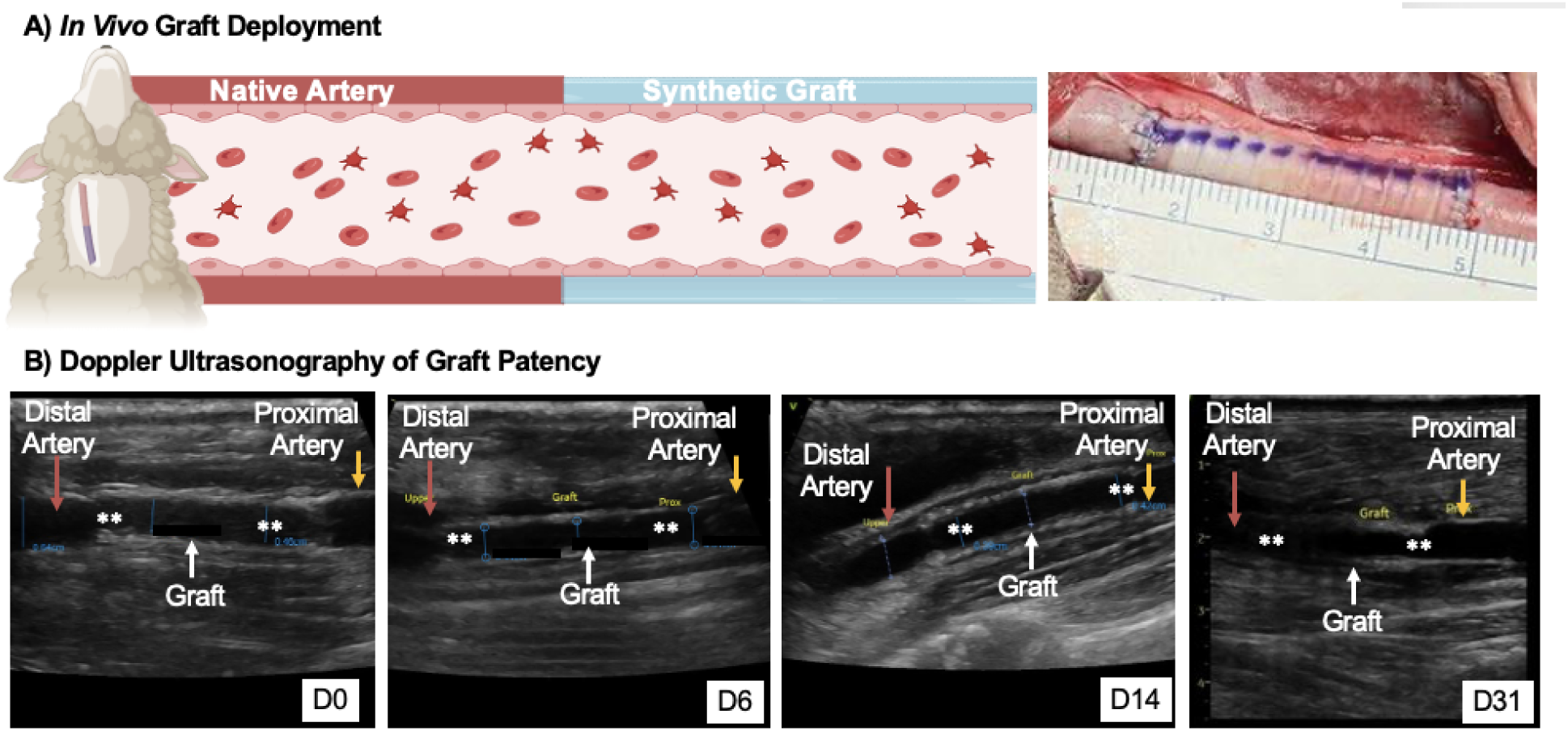
Pilot *in vivo* evaluation of multilayer vascular grafts. A) *In vivo* graft deployment in the carotid artery of an ovine model. B) Ultrasound at different time points of the implanted grafts (white arrow) joined to the proximal (orange arrow) and distal arteries (red arrow) at the anastomoses (white **) after immediate implantation and following 4 weeks of implantation confirming graft patency post-implantation and unobstructed blood flow through the graft.

## Discussion

There are currently no widely adopted synthetic small-diameter vascular grafts for CABG procedures due to high rates of complications from thrombosis.^26^ PEG-based hydrogels have the potential to improve thromboresistance of synthetic blood-contacting devices due to their innate antifouling properties and facile incorporation of bioactive cues to support adhesion, spreading, and migration of endothelial cells.^15, 16, 23, 27^ However, their conventional brittle nature and low extensibility make them prone to damage during surgical implantation. ^13, 21^ To this end, our lab developed a bioactive PEG-based interpenetrating network (IPN) hydrogel that couples high extensibility for damage resistance and integrin-binding motifs to support endothelialization post-implantation.^13^ This report describes the translation of this thromboresistant and damage-resistant IPN hydrogel to a bioactive coating for synthetic multilayer vascular grafts towards the development of off-the-shelf blood-contacting devices.

First, we investigated the difference between hydrogel physical properties and cell attachment when tested as a bulk hydrogel slab versus a coating of electrospun grafts. Our lab previously demonstrated that PEG-Scl2_GFPGER_ hydrogel substrates can facilitate α1β1 and α2β1 integrin engagement and that endothelial cell behavior can be modulated by changing ligand concentration, ligand identity, and gel stiffness.^23, 28, 29^ Scl2_GFPGER_ is a recombinant protein engineered to present integrin-binding motifs within a non-fouling PEG matrix that limits non-specific protein interactions.^30^ This design enables a high degree of selectivity in ligand presentation independent of substrate stiffness and physical properties, which is a key advantage in bioactive material systems.^30, 31^ In contrast to native ECM hydrogels, which contain multiple bioactive domains with variable receptor affinities and ligand concentration is often linked to other biophysical properties such as stiffness, the Scl2 platform provides a modular system for controlling endothelial cell–material interactions.^23, 29, 30, 32^ This enhanced control enables preferential engagement of α1β1 and α2β1 integrins and decoupling of gel properties (e.g., stiffness, ligand concentration).^23, 30^ In this study, we sought to validate that findings from these bulk hydrogel slab studies translate to hydrogel-coated grafts, confirming that these simple hydrogel substate studies are sufficient for optimizing graft performance and predicting endothelial cell behavior. This validation is critical as these hydrogel slab experiments enable precise isolation of substrate variables with opportunity for higher throughput, permitting systematic exploration of stiffness and ligand presentation before transitioning to more complex multilayer grafts. Despite the large difference in stiffness between the fiber meshes and hydrogel substrates, there was no observable differences in substrate modulus as measured by nanoindentation or protein incorporation. As expected, based on these findings, there was no difference in endothelial cell attachment and spreading on hydrogel slabs and hydrogel-coated meshes, suggesting that the endothelial cells primarily sense the hydrogel layer rather than the underlying mesh. Since endothelial cells are sensitive to variations in local substrate stiffness, a uniform hydrogel presentation of stiffness ensures consistent cell behavior and reduces heterogeneity in mechanotransduction pathways such as focal adhesion formation.^24, 33^ By confirming that the cellular response is solely due to the hydrogel coating, we are able to decouple the design criteria of each layer and independently optimize the hydrogel for biological performance and the electrospun mesh for biomechanical performance.^34^

Post-fabrication processing and sterilization can also impact device performance, particularly with regards to cellular responses to this bioactive hydrogel coating.^35^ The effect of sterilization on material properties and cellular responses for PEG-based substrates and other polymeric hydrogels have been previously characterized.^36–43^ Unlike other common sterilization methodologies, ethylene oxide sterilization was shown to have minimal effect on the PEG backbone when assessed through NMR analysis.^43^ Ethylene oxide sterilization performed on hydrogels was also previously reported to have no impact on surface roughness or radical concentration in comparison to untreated samples.^40^ Our lab has previously demonstrated that ethylene oxide sterilization had minimal effect on PEG-based hydrogels and subsequent cell-material interactions.^36, 44^ In this study, sterilization of our multilayer grafts showed no significant effect on hydrogel stiffness or endothelial cell attachment and spreading. Furthermore, there was no loss of damage resistance after drying and sterilization. Together, these studies indicate that sterilization does not significantly impair coating integrity or bioactivity retention. After only 3 hours of culture, the observed cell morphology and attachment indicate that the bioactive coating retained its ability to support early endothelial cell interactions following sterilization. Future studies evaluating longer culture durations under physiologically relevant flow conditions will be necessary to assess subsequent endothelial migration, proliferation, and formation of a confluent endothelial layer.

Resistance to coating failure during surgical handling remains a critical design requirement for hydrogel-coated vascular grafts, as hydrogel fragmentation or delamination may increase the risk of thrombosis or emboli. In the present study, dried and sterilized IPN-coated grafts maintained damage resistance following suturing, torquing, and tensile stretching, whereas PEGDA control coatings exhibited visible brittle failure. The reduced particulate generation and absence of macroscale coating fracture observed in the IPN formulations suggest the ability of the coating to withstand surgical manipulation and implantation procedures. The improved damage resistance observed in the IPN-coated grafts is supported by previously established tensile and fracture characterization of this hydrogel system.^13^ Prior studies demonstrated that PEUDAm IPNs exhibited substantially greater elongation at break and tensile strength than brittle PEGDA 3.4 kDa hydrogels, indicating that the network was capable of sustaining large strains without cohesive failure.^13^ Additionally, single-edge notch fracture testing demonstrated markedly higher maximum force at fracture in the IPN networks compared to single-network controls, supporting enhanced defect tolerance and mechanical energy dissipation behavior.^13^ The reduced particulate generation and absence of coating damage observed in the IPN formulations suggest the ability of the coating to withstand surgical manipulation and implantation procedures.

Following implantation, synthetic vascular implants undergo continuous pulsatile loading for the lifetime of the device. As such, it is important to assess the performance of the multilayer graft under these conditions prior to proceeding with *in vivo* testing. Our initial benchtop assessment demonstrated that the multilayer graft retained integrity and bioactivity throughout 6 weeks of physiological conditioning. This indicates the stability of the Scl2 protein presentation for at least 6 weeks under physiological loading conditions. This is consistent with previous reports of endothelial cells seeded on PEG hydrogels with Aam-PEG-I functionalized proteins under static conditions and highlights the hydrolytic stability of the Aam-PEG-I linker developed in our lab.^44^ Previous studies have also demonstrated that PEG-Scl2 hydrogels support maintenance of a thromboresistant endothelial phenotype, including regulation of pro-thrombotic markers (vWF, TF) and anti-thrombotic markers (eNOS, ADAMTS-13, tPA, and TFPI), further supporting the long-term functional bioactivity of these coatings. ^13, 29^ In addition to promoting endothelial adhesion and spreading, sustained presentation of these bioactive cues may help support long-term endothelial function during gradual transanastomotic endothelialization. Given that *in vivo* endothelial coverage progresses slowly from the anastomoses (∼0.2mm/day), prolonged bioactivity over long time frames is critical to confer a quiescent endothelial cell monolayer.^45^ The engineered graft presented here provides a promising method towards coupling long-term bioactivity and coating integrity with damage resistance.

To demonstrate suitability for *in vivo* graft testing, acute thromboresistance was first evaluated with static platelet attachment studies. Multilayer grafts demonstrated a significant reduction in platelet attachment in comparison to ePTFE (GORE-TEX®) vascular grafts under static conditions. Short-term whole blood flow studies were also used to simulate physiological conditions and further validate acute thromboresistance. Hydrogel-coated grafts demonstrated no visible signs of clot formation after 6 hours in comparison to ePTFE vascular grafts which had significant signs of platelet aggregation after 1-2 hours of exposure to whole blood. These results follow similar trends to previously reported platelet attachment and whole blood flow studies with bioactive PEG-based hydrogels, demonstrating enhanced thromboresistance as compared to ePTFE (GORE-TEX®).^13, 36^ The contrasting differences in substrate thrombogenicity highlights the ability of the multilayer grafts to maintain hemocompatibility and highlights the importance of evaluating the graft-blood interface for investigating acute thromboresistance prior to implantation. Although the current whole blood loop study focused on platelet attachment and visible clot formation, previous studies with PEG-based hydrogels incorporating Scl2 have also demonstrated minimal platelet activation, as measured through platelet activation assays and PAC-1 expression analyses.^29^ These findings suggest that the thromboresistant behavior of PEG-Scl2 hydrogels extends beyond reduced platelet attachment and includes mitigation of platelet activation pathways that could otherwise contribute to thrombosis. The acute thromboresistance in our graft is attributed to the antifouling hydrogel interface leading to reduced nonspecific protein adsorption and mitigation of subsequent platelet attachment.^15, 16, 46, 47^ As acute thromboresistance remains an early failure mechanism for blood-contacting devices, our graft overcomes a critical barrier to clinical translation.^48^

Synthetic vascular grafts often fail *in vivo* due to thrombotic occlusion, making *in vivo* assessments of thromboresistance and long-term patency essential for validating blood-contacting biomaterials.^48–50^ We selected an ovine model for this evaluation as they show the most similarity to humans in terms of the inflammatory response and thrombogenicity. They possess a coagulation system that is closer to the human than either dogs or pigs and have commonly been used as a model for intimal hyperplasia.^51–53^ Furthermore, this model permits implantation and monitoring of conduits as arterial grafts with sizing relevant to human anatomy (ID=4-6 mm) and catheter-based imaging with low luminal damage.^53, 54^ The ovine carotid artery is readily accessible, and carotid interposition grafts are well tolerated with minimal postoperative morbidity.^51^ In this study, coil-reinforced multilayer graft was utilized to mitigate kinking during long-term physiological assessments.^14^ For all cases, the grafts were successfully surgically implanted with good arterial size matching, stable anastomoses displaying minimal leakage, and maintaining pressure after hemostasis was established. Consistent with *in vitro* testing, all multilayer grafts implanted in an ovine carotid artery model displayed acute thromboresistance (6 hr); however, extended thromboresistance and patency was complicated by protocol iterations with only one animal proceeding for the full 30-day assessment. However, long-term patency was not achieved in all pilot animals, highlighting the complexity of translating in vitro biomaterial performance into the in vivo environment. Specifically, thrombosis observed in the first animal study prompted iterative refinement of postoperative anticoagulation and monitoring protocols, which contributed to improved outcomes in subsequent evaluations. These findings emphasize the importance of optimizing surgical, anticoagulation, and monitoring strategies when assessing novel blood-contacting materials in clinically relevant animal models. Although the in vitro hemocompatibility studies demonstrated reduced platelet adhesion under static and dynamic conditions, these assays do not fully capture the complex biological processes that contribute to in vivo thrombosis, including coagulation activation, inflammatory responses, protein adsorption, and interactions with circulating blood components over extended periods. Therefore, the observed differences between in vitro and in vivo performance highlight the necessity of clinically relevant animal models for comprehensive evaluation of vascular graft technologies. Despite these challenges, this initial pilot evaluation demonstrates the potential of the multilayer graft as an off-the-shelf small-diameter conduit for vascular interventions. Future studies will include additional replicates and incorporate comprehensive histological evaluation as a critical component of long-term graft assessment, including characterization of endothelialization through endothelial markers (e.g., CD31 and VE-cadherin), inflammatory cell infiltration, and neointimal remodeling. These analyses will be necessary to fully validate the biological effects of the hydrogel coating and determine its role in promoting endothelial coverage and preventing late-stage occlusion. Although the carotid interposition model provides a robust platform for initial evaluation of graft thromboresistance, handling, and patency, it does not fully replicate the complex hemodynamic environment associated with clinically relevant end-to-side bypass configurations, such as coronary artery bypass grafting (CABG).^55^ Therefore, future studies will investigate these multilayer grafts in end-to-side and CABG-relevant implantation models to further assess clinical translatability under physiologically representative flow conditions.

In comparison to currently investigated synthetic vascular graft platforms, the multilayer graft developed in this study provides several translational advantages that support its potential as an off-the-shelf blood-contacting device. Clinically used synthetic grafts (e.g., ePTFE, Dacron) continue to demonstrate poor patency in small-diameter applications due to thrombosis, compliance mismatch, and limited endothelialization.^56^ In contrast, the multilayer graft presented here provides two primary advantages over current synthetic grafts: 1) a reinforcing electrospun mesh with enhanced biomechanical matching to prevent intimal hyperplasia; 2) a bioactive and thromboresistant IPN hydrogel coating designed to simultaneously mitigate platelet attachment and support endothelial cell adhesion. Importantly, recent work from our group demonstrated that the same multilayer graft platform can be reproducibly manufactured using advanced manufacturing approaches, supporting scalability and manufacturability for clinical translation.^14^ Unlike many tissue-engineered vascular graft systems that require prolonged cell culture, patient-specific cell sourcing, or pre-endothelialization prior to implantation, this platform is designed as a fully synthetic construct used for post-implantation endothelialization with sustained bioactivity retention.^57, 58^ The modular graft design also enables independent optimization of the electrospun mesh for biomechanical performance and the hydrogel layer for hemocompatibility and endothelialization, providing greater flexibility than single-component graft systems.^59^ These design characteristics position the multilayer graft as a promising candidate for clinically translatable small-diameter vascular graft applications requiring durability, manufacturability, thromboresistance, and long-term endothelialization potential.

In this work, we developed and evaluated multilayer vascular grafts incorporating a durable, bioactive IPN hydrogel coating designed to address the primary failure modes of small-diameter synthetic conduits. By combining a high compliance electrospun polyurethane graft with a damage-resistant, endothelialization-promoting hydrogel coating, this platform integrates arterial biomechanical matching with biological functionality. By demonstrating that the hydrogel coating of the multilayer graft retained similar modulus and bioactivity as previous bulk hydrogel slab testing, we were able to confirm that optimization in 2D can be used to predict graft performance. In addition, the hydrogel coating showed no damage and maintained similar physical properties and bioactivity following ethylene oxide sterilization and prolonged physiological loading, supporting their suitability as off-the-shelf devices. Functional evaluation demonstrated sustained coating durability under surgical manipulation and dynamic loading, acute thromboresistance in a whole-blood bioreactor, and stable performance in a pilot ovine carotid interposition graft model. Together, these findings highlight the potential of these multilayer vascular grafts with durable bioactive coatings as functional, off-the-shelf small-diameter conduits for bypass and other vascular procedures. In the future, additional replicates will be included in the ovine model to establish consistency in long-term patency assessments and investigate endothelialization on the synthetic vascular graft constructs.

## Methods Materials

All chemicals were purchased from Sigma-Aldrich and used as received unless otherwise noted.

### Synthesis of polyether urethane diacrylamide (PEUDAm)

PEUDAm was synthesized in a three-step process as previously reported.^13^ Briefly, PEG (20 kDa, 1 equiv., 1.74mmol) was dissolved in DCM and reacted with CDI (15 equiv, 26.1 mmol) purged with nitrogen for 2 h at 30-35C. The reaction continued mixing overnight prior to quenching with water and solvent removal to collect PEG-CDI. The product was washed with deionized water three times in a separatory flask, dried with sodium sulfate, and collected via vacuum filtration. Solvent was removed under vacuum with a final drying under vacuum at 65°C for 1 hour. PEG-CDI was then added dropwise addition to EDA (15 equiv.) at room temperature under nitrogen atmosphere and the mixture stirred overnight. The PEG-EDA product was washed with deionized water three times in a separatory flask, dried with sodium sulfate, and collected via vacuum filtration. The reaction was precipitated in 10× volume of ice-cold ether, collected via vacuum filtration, and dried under vacuum at 65 °C for 3 h. PEUDAm was then synthesized by functionalizing the PEG-EDA product with acrylamide groups through the addition of triethylamine (4 equiv) and acryloyl chloride (4 equiv) to PEG-EDA (1 equiv). The reaction was stirred for 48 h then quenched using potassium carbonate. The product was washed with water two times, dried with sodium sulfate, and collected via vacuum filtration. Precipitation was run in ice-cold diethyl ether and collected via vacuum filtration. The final product was dried at atmospheric pressure overnight, then briefly under vacuum. The chemical structure of the final PEUDAm product was verified with proton nuclear magnetic resonance H^1^ NMR (CDCl_3_): δ = 6.94 (broad s, 2H, –C–N*H*–CH_2_–), 6.28 (dd, 2H, *H*_2_C CH–C–), 6.17 (m, 2H, H_2_C C*H*–C–), 5.85 (broad s, 2H, –C–N*H*–CH_2_–), 5.61 (dd, 2H, *H*_2_C CH–C–), 4.20 (m, 4H, –H_2_C–C*H*_2_–O–), 3.65 (m, 1720H, –O–C*H*_2_–CH_2_–), 3.34 (m, 4H, –CH_2_–C*H*_2_–NH–). Polymers with percent conversions of hydroxyl to acrylamide end groups over 80% were used in this work.

### Fabrication of electrospun grafts

Electrospun grafts were fabricated for benchtop testing of multilayer grafts. An 18wt% solution of Bionate^TM^ 80A in a solvent mixture of N,N-dimethylacetamide and tetrahydrofuran was pumped at 0.5 ml/h through a 20-gauge blunt needle onto a rotating 5 mm-diameter mandrel (500 RPM). To enable facile removal of the graft, the 5 mm rod was dip-coated in a 5% 35 kDa PEG in dichloromethane prior to fiber collection. The mandrel and needle were set at a 50-cm working distance and charged to -5 kV and +17-20 kV, respectively. Fibers were collected for 1 hour under ambient conditions (45–50%, temperature 21–22 °C). After spinning, coated rods were immersed in water for at least 1 hour to dissolve the PEG layer for graft removal and then grafts were dried under vacuum overnight at < 0.1 mbar (1400E Benchtop Vacuum Oven, VWR, Radnor, PA). Fiber diameter was analyzed via scanning electron microscopy (Phenom Pro, NanoScience Instruments, Phoenix, AZ,). Samples were coated with 5 nm of gold (Sputter Coater 108, Cressington Scientific Instruments, Hertfordshire, UK) and imaged at 5000X magnification. Average fiber diameter was characterized in ImageJ by drawing a mid-line and measuring the diameter of the first ten fibers that crossed the line (*n* = 5 images per graft).

Coil-reinforced electrospun grafts were utilized for *in vivo* evaluation to ensure sufficient kink resistance for surgical implantation and testing. Briefly, coil-reinforced grafts were fabricated in three layers using a combination of electrospinning and 3D printing as described in Robinson et al.^14^ Briefly, the first electrospun layer was fabricated as described above with an initial mesh thickness of ∼120–150 µm was targeted for the first layer. A polymeric coil was then applied to the grafts via solution deposition with a custom coil printer. A coil with spacing of 2.5 mm and a diameter of 0.35 mm was applied using a Bionate^TM^ 80A solution (21.5wt % in 1,1,1,3,3,3-Hexafluoro-2-propanol). The coil-reinforced graft was then dried and a second electrospun layer was applied with 18wt% Bionate^TM^ 80A solution as described above. Grafts were allowed to dry overnight and then removed by placing the mandrel in water for ∼30 min to dissolve the PEG layer and sliding the graft off. Grafts were trimmed to ∼4 cm prior to characterization. To improve integrity between the electrospun and coil layers, coil-reinforced grafts underwent solvent vapor welding. Briefly, a Teflon rod (5 mm in diameter, 100 mm long) was placed inside the grafts and the grafts exposed for two hours in a dessicator with a 4o mL of tetrahydrofuran. Following solvent vapor fusion, the grafts were dried under vacuum overnight at < 0.1 mbar (1400E Benchtop Vacuum Oven, VWR, Radnor, PA).

### Bulk Hydrogel Slab Fabrication

IPN hydrogels were fabricated as previously described.^13^ First network hydrogels were fabricated with 10 wt/vol% PEUDAm 20 kDa macromer solutions. Irgacure 2959 (10 wt/vol% solution in 70 vol% ethanol) was added at a final concentration of 0.1 wt/vol% to hydrogel precursor solutions. Precursor solutions were then pipetted between 0.5 mm spaced glass plates and cured on a UV transilluminator (UVP, 25 watt, 365 nm) to form bulk first network hydrogels. Hydrogels were then swollen to remove residual macromer, then dried at ambient pressure overnight. Dried hydrogels were placed in a vial with a solution of N-acryloyl glycinamide (NAGA; BDL Pharma) (20 wt/vol%), bis-acrylamide (0.5 mol% relative to NAGA), and Irgacure 2959 (2 wt/vol%). Soaking solutions were rotated overnight and protected from light. IPNs were cured by placing monomer-swollen hydrogels between glass spacer plates set to the swollen thickness and curing on a UV transilluminator for 20 minutes on both sides.

### Multilayer graft fabrication and characterization

A diffusion-mediated redox initiation and hydrogel crosslinking was utilized to coat the vascular graft with the IPN hydrogel as previously described.^22^ Briefly, grafts were soaked with a graded ethanol/water ramp (70 vol%, 50 vol%, 30 vol%, and 0 vol%, 15 minutes each) to promote wetting and aqueous solution penetration prior to coating. Grafts were immersed in a solution of 3 wt% iron gluconate (IG) in deionized water for 15 minutes. Following a brief dip in methanol, mesh substrates were submerged in an aqueous solution of 10 wt/vol% PEUDAm 20 kDa with ammonium persulfate (APS, 0.05 wt/vol %) and 2 wt/vol % Irgacure 2959 for 20 s. After dip coating, grafts were set on a UV plate (Intelligent Ray Shuttered UV Flood Light, Integrated Dispensing Solutions, Inc., 365 nm, 4 mW/cm^2^) for 12 min for additional photoinitiated crosslinking of the PEUDAm hydrogel layer followed by overnight drying. After drying, grafts were immersed in DI water for 5 h with water exchanges done after 10 min, 1 hour, and 3 h to remove unreacted reagents. Substrates were then dried overnight and then soaked overnight in a second network precursor solution comprised of 20 wt/vol % N-acryloyl glycinamide (NAGA, BLD Pharma), 0.25 mol % bisacrylamide (bisAAm), and 2 wt/vol % Irgacure 2959 overnight protected from light at 4 °C. Prior to photocuring, functionalized Scl2_GFPGER_ (functionalization of ∼10% of the available lysines with acrylamide-PEG-isocyanate) was added to the lumen of the NAGA-swollen composites.^28^ Grafts were then set on a 4 mm glass rod and placed on a custom rotating device positioned on a UV transilluminator to initiate UV cure of the IPN hydrogel. The resulting multilayer grafts were then dried overnight to facilitate additional crosslinking, followed by soaking in water to remove unreacted reagents. Multilayer grafts were then either tested immediately or sterilized via ethylene oxide prior to testing.

Multilayer grafts with the brittle PEG-diacrylate hydrogel coating were fabricated as a positive control for surgical damage.^13, 21^ Briefly, electrospun mesh sleeves were taken through a graded ethanol/water soak and then soaked in a precursor solution of 10 wt/vol% PEGDA 3.4 kDa. Electrospun meshes were then placed in a cylindrical mold with an inner glass mandrel (3 mm OD) and the precursor solution was added to the mold and crosslinked with UV light for 12 min in a custom-built UV box. The resulting multilayer grafts were then dried overnight to facilitate additional crosslinking, followed by soaking in water to remove unreacted reagents.

### Nanoindentation characterization of hydrogel coating modulus

Nanoindentation experiments were performed with the Piuma Nanoindentor (Optics11 Life, Amsterdam, Netherlands). Data was collected using the Piuma software V3.4.7. Modulus measurements were assessed at an indentation depth of 2 µm with indentation speeds of 2 µm/s. Moduli values were assessed a 2×2 matrices 1000 µm apart on three graft sections (n = 3 grafts per condition, 4 indentations per graft). The data from the Piuma software was fit to the Hertzian model and a python script was used to validate the data.

### Characterization of hydrogel damage resistance

The effect of drying and sterilization on the damage-resistance of the IPN hydrogel coating was assessed in comparison to a brittle PEG diacrylate (3.4 kDa) hydrogel positive control.^13, 21^ Suture-induced damage was assessed as previously described by Post *et al.*^21^ Briefly, a 7-0 Ethicon suture was passed through the hydrogel substrates (n = 3 grafts, 4 passages per graft). Resulting particulates were then visualized under a stereoscope (Olympus SZ61), and the number of particulates dislodged were counted manually. Torque-induced damage was evaluated as previously described by Wancura *et al*.^13^ Briefly, tubular composites were gripped at the center region with forceps and twisted 180° (n = 3 grafts). Resulting damage to the inner layer was then visualized under a stereoscope (Olympus SZ61). Hydrogel delamination under uniaxial tensile loading was assessed as an additional measure of coating durability.^13^ Briefly, multilayer grafts were sectioned into 20 x 5 mm dimensions and dyed with food coloring to visualize hydrogel coating damage (n = 3 grafts, 3 samples per graft). Specimens were mounted between the clamps of a Dynamic Mechanical analyzer (TA Instruments) and stretched at a rate of 1 mm/s to 100% strain under video recording to visualize the onset of coating damage.

### Physiological conditioning of multilayer grafts

A pulsatile flow setup was utilized to characterize the effect of physiological loading on coating integrity and bioactivity retention of multilayer grafts. A physiological loading setup was designed with a Masterflex L/S variable speed peristaltic pump (Cole-Parmer) fitted with a Masterflex L/S pump head to provide pulsatile flow and promote pressure at intraluminal pressure. Multilayer grafts were mounted to a custom testing chamber and underwent pulsatile loading for 24 hours, 1 week, and 6 weeks with exposure to 120/80 mmHg at 37°C to determine (n = 3 grafts per time point). Delamination was evaluated at the graft edges and along the center via visual assessment of the graft. Specimens were punched along the length of the conditioned grafts (8 mm diameter, n = 3 specimens per graft) and analyzed for protein retention and bioactivity via endothelial cell adhesion and spreading.

### Protein incorporation and retention

Bioactive hydrogels and multilayer vascular graft sections were analyzed for Scl2 protein content as described previously.^28^ Specimens (8 mm diameter) were first treated in a solution of 3% hydrogen peroxide with 1.25 mM cobalt chloride for accelerated oxidative degradation to release incorporated protein (n = 9 bulk hydrogel slab specimens; n = 3 grafts per condition with 3 specimens per graft). The solutions were then diluted in DI water, frozen, and lyophilized to isolate hydrogel components. The lyophilized powder was then resuspended in PBS, and the protein content was evaluated with the 3-(4-carboxy-benzoyl)-2-quinoline-carboxaldehyde (CBQCA) protein quantification kit (Molecular Probes, Life Technologies) according to the manufacturer protocol. Briefly, 10 µl sample aliquots were diluted in 125 µl of 0.1 M sodium borate. Then, 5 µl of 5 mM potassium cyanide (KCN) was added along with 10 µl of 5 mM CBQCA. Following a 1-3 h incubation with shaking, fluorescence of all samples and protein standards were assessed in triplicate with an Infinite M200 Pro plate reader (excitation = 465 nm, emission = 550 nm). Standard curves of fluorescence were generated with known protein concentrations.

### Endothelial cell adhesion and spreading

Bioactivity retention was assessed via endothelial attachment and spreading assessments on bulk hydrogel slabs and multilayer grafts before and after physiological loading for up to 6 weeks. Human thoracic aortic endothelial cells (HTAECs) were cultured in ECGM-2 cell media (Promocell) and seeded at 10,000 cells (8 mm diameter punched specimens; n = 3 grafts per condition, 3 specimens per graft). Cells attachment and spreading was evaluated after 3 hours. Hydrogel and multilayer graft specimens were fixed with 3.7% glutaraldehyde and stained with rhodamine phalloidin (actin/cytoskeleton) and SYBR green (DNA/nucleus). Specimens were imaged with a fluorescence microscope (Nikon Eclipse TS100) and cell adhesion and spreading were quantified as previously described.^60^

### Thromboresistance assessment of multilayer constructs

As an initial assessment of graft thromboresistance, platelet attachment to multilayer grafts was compared to an ePTFE clinical control graft as described previously.^13^ Briefly, multilayer graft and ePTFE vascular graft (GORE-TEX®, Gore Medical) specimens (8 mm, *n* = 12) were soaked in fetal bovine serum overnight at 37 °C, washed once with PBS to remove non-adhered protein, and placed in a 48 well plate. Platelets were isolated from human whole blood drawn from a volunteer as described previously.^13, 29^ Informed written consent was obtained from the volunteer as described in the approved IRB protocol STUDY00002311. Platelets were resuspended in Tyrode’s buffer at half the original volume of the platelet rich plasma and stained with Sudan B Black solution (5% in 70% ethanol) at a 1:10 ratio for 30 minutes at room temperature. A platelet suspension (500 μL, 10 × 10^6^ platelets per mL) was added to each graft specimen, and platelets were allowed to adhere to substrate for 30 min at 37 °C on a shaking incubator. Samples were gently washed twice to remove unattached platelets and then placed in 150 μL DMSO for 15 min at room temperature. The lysate was then diluted with 150 μL PBS and solutions read in triplicate on a spectrophotometer (Tecan Infinite M Nano+, absorbance measured at 616 nm).

A dynamic whole blood reactor was then used to assess full size vascular grafts as a more physiologically relevant measure of thromboresistance. Briefly, heparinized whole blood was flowed through the lumen of multilayer grafts for 6 hours and platelet attachment assessed. A 6 mm internal diameter ePTFE vascular graft served as a clinical control (GORE-TEX®, Gore Medical). Fresh, heparinized whole blood was incubated with mepacrine (10 μM) for 30 min prior to testing fluorescently label platelets. The grafts were fitted onto custom chambers connected to a peristaltic pump (Masterflex, Cole Parmer) and blood flowed through the lumen at 37 °C with a volumetric flow rate of 120 ml/min for 6 hours (n = 3 grafts). After 6 hours of constant flow, grafts were removed, gently rinsed with PBS, and fixed in formalin (10% v/v) overnight at 4 °C and imaged with a fluorescence microscope (Nikon Eclipse TS100) in 3 randomly selected fields of view per specimen to analyze platelet adhesion.

### Ovine carotid artery interposition graft model

All animal procedures were approved by the Institutional Animal Care and Use Committee (IACUC) at Baylor College of Medicine and performed in accordance with institutional and NIH guidelines for the care and use of laboratory animals. Sheep underwent preoperative imaging to estimate carotid artery diameter for size-matching with coil-reinforced multilayer grafts. Perioperative antithrombotic management included systemic heparinization prior to vascular clamping and graft implantation. After neck cutdown and isolation of the carotid artery, end-to-end anastomosis was performed as an interposition graft using continuous 7-0 prolene sutures into the left carotid artery (n = 3 grafts). Postoperative antithrombotic management included antiplatelet therapy and systemic anticoagulation, including combinations of aspirin, clopidogrel, intravenous heparin, and oral warfarin, with serial coagulation monitoring (including ACT and PT/INR assessments) performed during postoperative follow-up. These postoperative anticoagulation and surveillance protocols were iteratively refined across the three cases. In the first case, anticoagulation consisted of an aspirin (325 mg) and clopidogrel (150-300 mg) loading dose. This was followed by continuous heparin infusion (100 U/ml) and with warfarin (3–6 mg) beginning on postoperative day 2, with Doppler ultrasound surveillance performed on a weekly interval. In the second case, heparin infusion (100 U/ml) was initiated immediately postoperatively with activated clotting time (ACT)-guided escalation (10-20 ml/hr) and multiple IV boluses (5000 U). This was supplemented with aspirin (650 mg) and warfarin (3 mg) on postoperative days 1–2. In the third case, this ACT-guided heparin and frequent coagulation monitoring were continued over a prolonged postoperative course with prolonged heparin infusion (100 U/ml), intermittent IV boluses (2000-5000 U) with infusion rate adjustments (up to 20 ml/hr). In addition, warfarin dosing was progressively increased (3–25 mg) to maintain therapeutic anticoagulation, and Doppler ultrasound surveillance was performed throughout the full 31-day observation period rather than at widely spaced intervals. Doppler ultrasonography was used to confirm graft patency and screen for intraluminal thrombus using a clinical ultrasound system (GE Vivid S70N, GE Healthcare) equipped with a high-frequency linear array transducer (9L-D probe). Animals were imaged in the supine position under anesthesia, and care was taken to avoid external compression of the graft during scanning. Color Doppler and pulsed-wave spectral Doppler were obtained using consistent acquisition practices across examinations. The Doppler cursor angle was aligned as parallel as possible to the direction of blood flow and maintained at ≤60°. Pulse repetition frequency and wall filter settings were adjusted as needed to optimize visualization of flow within the graft lumen. The pulsed Doppler sample volume (approximately 1–2 mm) was positioned at the center of the vessel lumen. For longitudinal consistency, Doppler assessments were performed at three predefined locations: (1) the native artery proximal to the anastomosis (approximately 0.5–1 cm upstream), (2) the mid-segment of the vascular graft, and (3) the native artery distal to the anastomosis (approximately 0.5–1 cm downstream). Vessel diameter was measured in B-mode at corresponding locations for anatomical reference. Patency and thrombus assessments were determined using prespecified Doppler criteria. Grafts were assessed macroscopically in situ before explantation for any signs of infection and anastomotic disruption. Visual inspection of the hydrogel coating for signs of delamination or degradation was also performed following explantation.

### Statistical analysis

The data for all measurements are displayed as mean ± standard deviation. An analysis of variation (ANOVA) comparison utilizing Tukey’s multiple comparison test was used to analyze the significance of data among multiple compositions. All tests were carried out at a 95% confidence interval (p < 0.05).

## Acknowledgment

This work was supported by the Roderick D. MacDonald Research Fund Award 24RDM002, National Institutes of Health grant number R01 HL180615, and the National Science Foundation Graduate Research Fellowship Program under Grant (2023355666). Any opinions, findings, and conclusions or recommendations expressed in this material are those of the author(s) and do not necessarily reflect the views of the National Science Foundation. Bionate^TM^ 80A was provided by the Biomedical division of dsm-firmenich (Exton, PA). The graphical abstract and cell-material renderings in Figures 1-4 and Figure 6 were created in BioRender.

## Author Contributions Statement

A.N: Writing – original draft, Methodology, Investigation, Formal analysis, Data curation, Conceptualization. A.F: Methodology, Investigation. N.A: Methodology, Investigation. M.L.: Methodology. A.R: Methodology. N.G.: Methodology, Investigation. X.Z.: Methodology, Investigation, Formal analysis, Data curation. LJ.G.: Methodology, Investigation, Formal analysis, Data curation. M.T.Z.N.S.: Methodology, Investigation, Formal analysis, Data curation. A.E.: Methodology, Investigation, Formal analysis, Data curation. E.C.H.: Supervision, Writing – review & editing, Conceptualization. All authors reviewed the manuscript

## Data availability

Data is available on Texas Data Repository at https://doi.org/10.18738/T8/N0EXFM.

