## Supplemental Information for "Off-the-Shelf Multilayer Vascular Grafts with Damage-Resistant Hydrogel Coatings Incorporating Integrin Targeting"

**Supplementary Data**

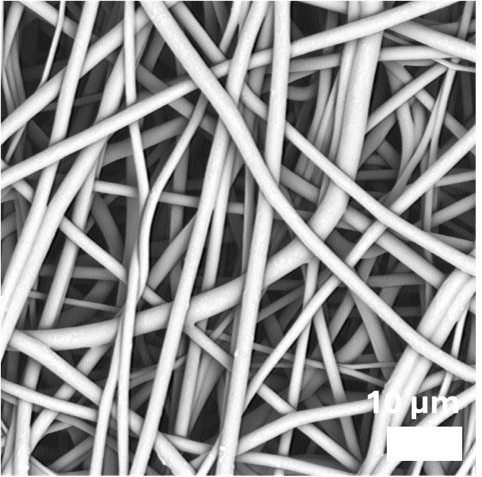

**Supplementary Figure 1**: Representative scanning electron micrograph of electrospun Bionate^®^ segmented polyurethane mesh with average fiber diameters 1.8 ± 0.5 µm (n = 3 grafts per condition).

**Supplementary Table 1.** Biomechanical properties of coil-reinforced multilayer vascular grafts (n = 3 grafts).

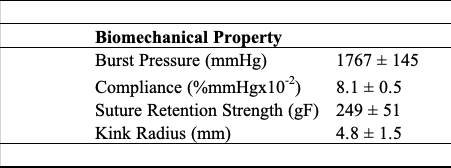

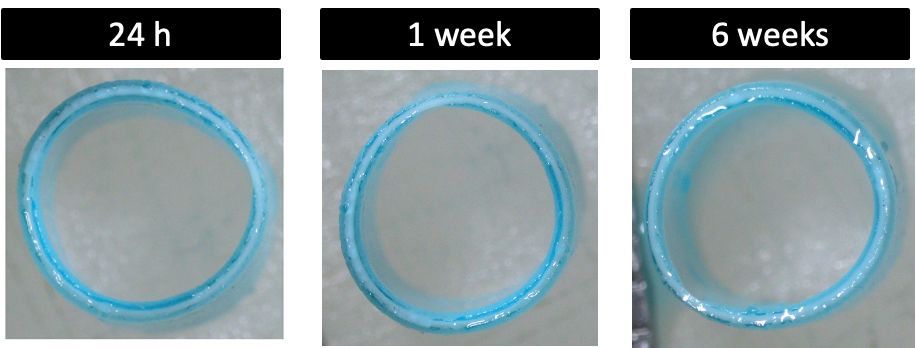

**Supplementary Figure 2:** Representative graft images following physiological loading for 24 h, 1 week, or 6 weeks (n = 3 grafts per timepoint).

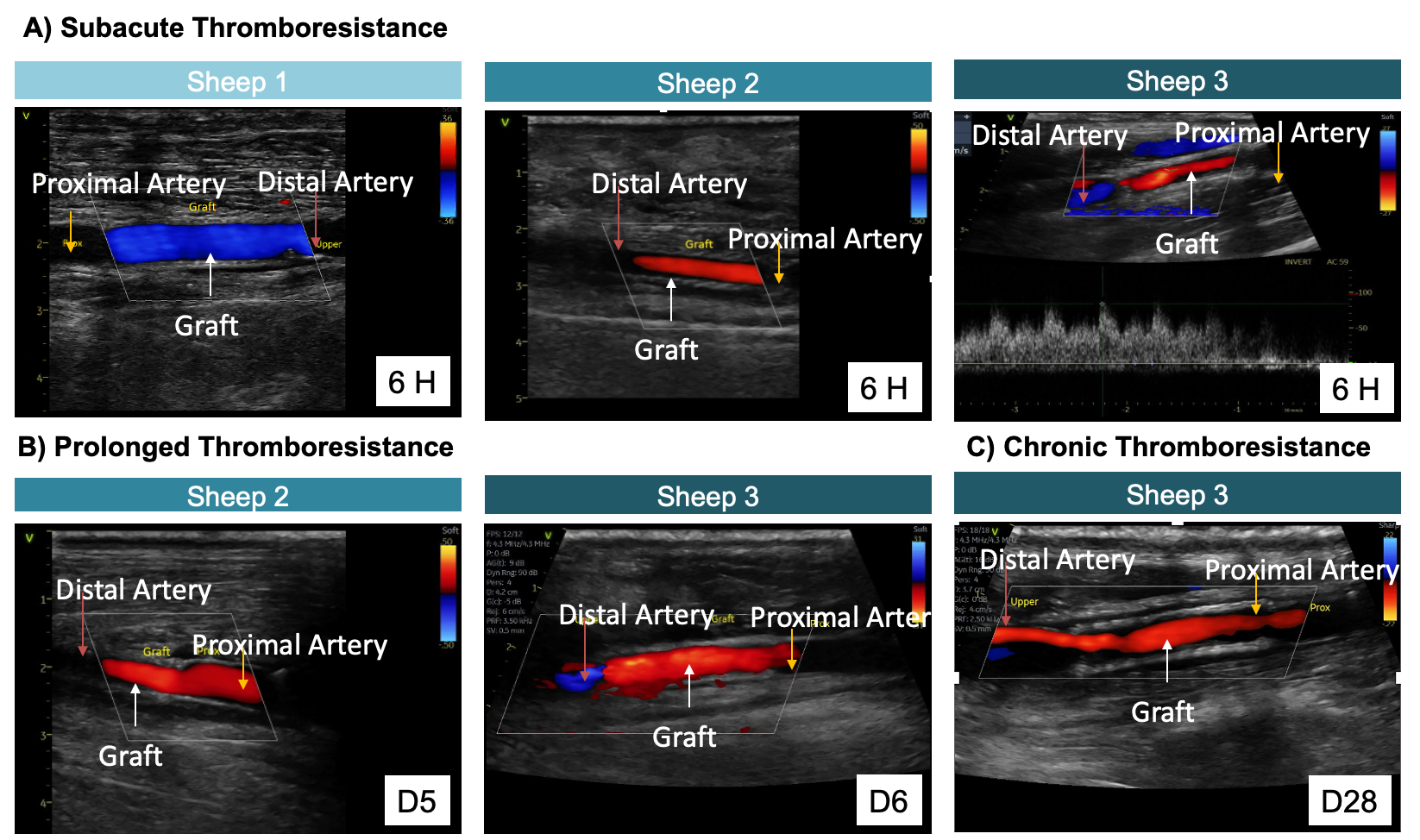

**Supplementary Figure 3:** Pilot in vivo evaluation of multilayer vascular grafts. Doppler ultrasonography showing A) subacute thromboresistance (6 h) for all three sheep B) prolonged thromboresistance (5-7 days) for two sheep, and C) sustained thromboresistance (28 days) for the third sheep. Images show grafts (white arrow) joined to the proximal (orange arrow) and distal arteries (red arrow).
